# Foraging behavior and patch size distribution jointly determine population dynamics in fragmented landscapes

**DOI:** 10.1101/2021.11.10.468021

**Authors:** Johannes Nauta, Pieter Simoens, Yara Khaluf, Ricardo Martinez-Garcia

## Abstract

Increased fragmentation caused by habitat loss represents a major threat to the persistence of animal populations. How fragmentation affects populations depends on the rate at which individuals move between spatially separated patches. Whereas negative effects of habitat loss on biodiversity are well-known, effects of fragmentation *per se* on population dynamics and ecosystem stability remain less understood. Here, we use a spatially explicit predator-prey model to investigate how the interplay between fragmentation and optimal foraging behavior affects predator-prey interactions and, subsequently, ecosystem stability. We study systems wherein prey occupies isolated patches and are consumed by predators that disperse following Lévy random walks. Our results show that the Lévy exponent and the degree of fragmentation jointly determine coexistence probabilities. In highly fragmented landscapes, Brownian and ballistic predators go extinct and only scale-free predators can coexist with prey. Furthermore, our results confirm that predation causes irreversible habitat loss in fragmented landscapes due to overexploitation of smaller patches of prey. Moreover, we show that predator dispersal can reduce, but not prevent nor minimize, the amount of lost habitat. Our results suggest that integrating optimal foraging theory into population- and landscape ecology is crucial to assessing the impact of fragmentation on biodiversity and ecosystem stability.

---

Loss of habitat presents a major threat to global biodiversity (1) and typically leads to fragmented landscapes that contain smaller and more spatially isolated patches in which local extinctions are more likely to occur (2, 3). While ecologists agree that habitat destruction, and the subsequent increase in habitat fragmentation, affects biodiversity negatively (4), the potential effects of fragmentation *per se* on population densities and species’ persistence are much less understood (5, 6). As it is known that fragmentation *per se* induces changes in demographic rates and drifts in population genetics (7), it is critical to assess its effects on population dynamics and ecosystem stability.

Fragmentation *per se* (hereafter; fragmentation) describes changes in the spatial habitat configuration without significant habitat loss (8). Theoretical and experimental studies indicate that fragmentation can result in larger species’ extinction probabilities, as small patches sustaining small populations are more sensitive to demographic fluctuations (8). In contrast, fragmentation might also favor species’ persistence by increasing immigration rates, patch connectivity and the diversity of habitat available within a smaller area (for a review, see Ref. 9). Therefore, whether, and how, fragmentation impacts species’ persistence strongly depends on the spatial configuration of the landscape (10, 11) and the dispersal behavior (i.e., movement between fragments) of individual organisms (12, 13). Despite the interplay between landscape structure and individual movement being essential for understanding population dynamics, research on each of these fields has progressed mostly independent from one another (14, 15). As a result, a general framework to investigate how fragmentation, movement behavior, and demographic rates jointly determine species’ persistence is lacking.

On the one hand, studies on individual movement are often grounded in optimal foraging theory (16). These studies investigate foraging behavior on short time scales and most often neglect demographic events and evolutionary processes (but see Ref. 17). Instead, they examine how individual movement behavior defines search times and study correlations between foraging efficiency and resource density (18). Often, movement is modeled using scale-free random searches, known as *Lévy walks*, in which displacement lengths are sampled from power laws with varying exponent (19). This particular choice for the distribution of displacement lengths is based on empirical observations reporting scale-free patterns in the movement of different species (20–24). Although these patterns in displacement lengths might be recapitulated both by memoryless Lévy searchers and area-restricted foragers (25, 26), we consider here the former because they provide a simple mathematical framework to explore how individuals balance the exploration-exploitation trade-off underlying search processes (27). In general, Lévy walks are a very efficient random foraging strategy in sparse resource landscapes (28–31), including fragmented landscapes (32, 33).

On the other hand, studies on population dynamics consider longer time scales and often assume simplified individual movement (34–37). Few studies have integrated optimal foraging behavior in population-based models (14, 38) and, to the best of our knowledge, only Ref. 39 studies population dynamics in a system of optimal foragers while considering fragmented prey habitat. The study, however, mostly controlled for habitat availability and did not systematically investigate how the complex spatial distributions of habitat as observed in natural landscapes impact population dynamics. Here, we extend the framework proposed in Ref. 39 and scrutinize these effects using techniques from landscape ecology that allow us to generate lattices with precise levels of fragmentation (40, 41).

To study the interplay between optimal forager movement, fragmentation, and demographic rates, we develop a stochastic, spatially explicit predator-prey model in fragmented landscapes. Fragmentation restricts prey individuals to inhabit spatially separated fragments, whereas predators are assumed to display natural (optimized) foraging behavior and disperse following a Lévy walk. By varying habitat fragmentation and predator movement, we quantitatively examine the effects of dispersal on ecosystem stability in fragmented landscapes.

## Stochastic predator-prey model in fragmented landscapes

We develop a stochastic predator-prey model in a twodimensional landscape with fragmented prey habitat. The landscape is represented by a periodic square lattice in which a fraction *ρ* ∈ [0,1] of the sites provide prey habitat. To investigate how predator movement and the spatial distribution of prey habitat jointly determine predator-prey population dynamics, we fix the fraction of prey habitat *ρ* and vary only the statistical properties of the patch sizes. In other words, we focus on the spatial configuration of habitat (fragmentation *per se*) and do not consider potential effects of habitat loss. The degree of fragmentation is determined by the spatial correlations in the distribution of prey habitat. In our model, these are controlled by the Hurst exponent *H* ∈ (0,1). In general, the limit *H* → 1 defines low habitat fragmentation, whereas *H* → 0 defines highly fragmented landscapes (Fig. 1 and see SI for more details).

**Fig. 1.**
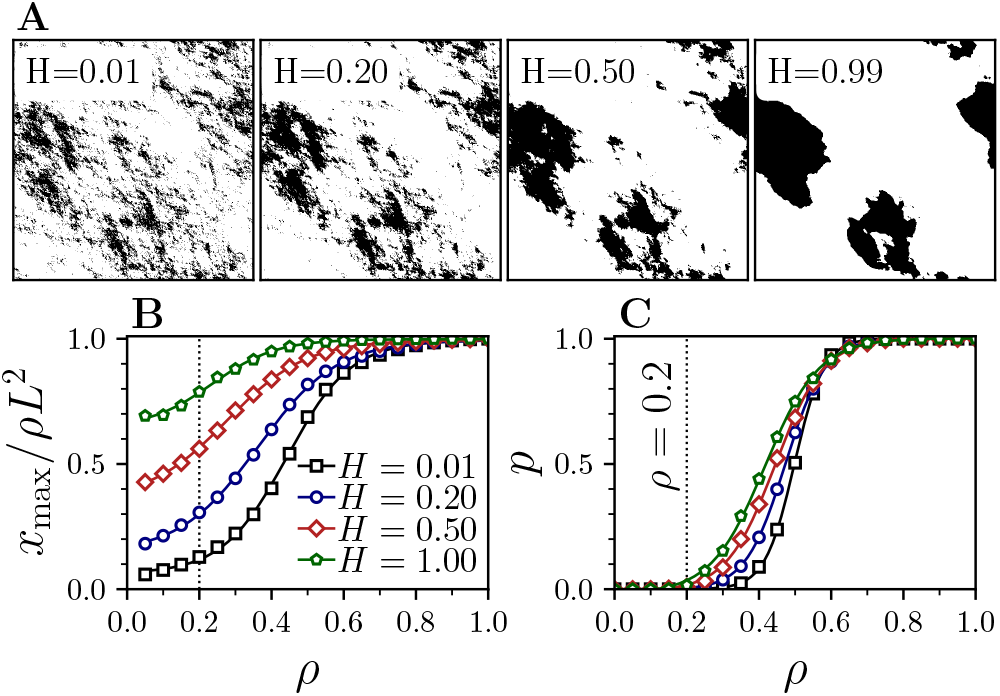
(A) Fragmented landscapes of prey habitat used in our model implementation with *L* = 512 and *ρ* = 0.2. The degree of fragmentation increases with the Hurst exponent, *H*. Black and white regions depict prey habitat and matrix, respectively. (B) Normalized maximum patch size *x*_max_ for different *H* versus habitat density *ρ* (see SI). (C) Percolation probability *p* as a function of *ρ* (see SI). Dotted vertical line indicates the habitat density *ρ* = 0.2 below the percolation threshold (*p* ≈ 0) used in our experiments.

We assume that individual prey are sessile, can only occupy habitat patches, and cannot survive in the matrix. Each time step, they reproduce with probability *σ* and can potentially die up encountering a predator. We assume predators are, in contrast, highly motile and perform Lévy walks in which the dispersal length *ℓ* follows a discrete power-law distribution *p*(*ℓ*) ∝ ℓ^-*α*^ with exponent 1 ≤ *α* ≤ 3. For *α* ≥ 3, predator movement converges to Brownian motion whereas the limit *α* → 1 recovers ballistic motion. In contrast to Lévy flights (which were discussed in Ref. 39), in which relocations are instantaneous, Lévy walks represent movement at constant velocity. Hence, in our model, individual predators move a fixed distance every time step (the unit lattice spacing) and the thus duration of each relocation event is proportional to the length of the displacement (19). Predator relocations can be interrupted by predator death or by an encounter with prey or other predators. When a relocation is interrupted, a new direction and dispersal length are sampled and the predator resumes its movement in the next time step.

For predator-prey encounters, we consider that when predators cross a site occupied by prey, the probability that they interact 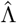 decays with the current dispersal length, i.e., 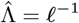. This assumption models intermittent search behavior (27), which combines phases of non-reactive long, straight displacements with reactive phases featuring shorter displacements and more frequent turns (42–44). In each predator-prey encounter event, prey is consumed and replaced with a new predator with probability *λ* (predator reproduction probability) and with the focal predator otherwise (no predator reproduction). See SI for further details on the parameters and the model implementation.

## Results

We simulate the predator-prey model on a square lattice of lateral length *L* = 512, with prey habitat density *ρ* = 0.2 and different levels of fragmentation 0 < *H* < 1. We choose *ρ* considering that fragmentation impacts landscape properties more strongly when habitat is not abundant (e.g., Refs. 1, 45, 46, see SI for more details) and that Lévy foraging maximizes prey intake only for low prey habitat density (e.g., Ref. 28). We define the spatially averaged predator and prey densities *N* and *M* and initialize our simulations with *M*_0_ = *N*_0_ = *ρ*. Predators are distributed randomly on the matrix and prey individuals fully occupy the habitat patches. Measurements of species densities were taken when the system converged to a quasi-stationary stable state after *T* = 10^4^ Monte Carlo time steps (Fig. SI.4). Results in this quasi-stationary stable state did not depend on the specific initial condition chosen for the simulations.

We are interested in investigating the impact of fragmentation in *fragile* ecosystems (cf. Ref. 39). That is, systems that are already close to a extinction threshold when habitat patches are large (*H* → 1). Hence, we parameterize demographic rates such that they bring the predator-prey dynamics close to an extinction transition (Ref. 39 and see SI) and consider fragmentation, defined by the Hurst exponent *H*, and predator dispersal, defined by the Lévy exponent *α*, as the only free model parameters. Using this simulation setup, we study the impact of habitat structure and predator dispersal on population dynamics, ecosystem stability and patterns of irreversible habitat loss.

### Population densities and species richness

We measure population sizes in the quasi-stationary stable state for different degrees of habitat fragmentation and foraging strategies. Since prey reproduction rate is fixed in our simulations, equilibrium population sizes are determined by predator-prey encounter rates and predator reproduction rates. The long-time prey population size decreases monotonically as predator movement goes from ballistic to Brownian (Fig. 2). Predator density, however, is maximal for an intermediate value of the Lévy exponent and its optimal value depends on the degree of fragmentation. For each degree of fragmentation *H* we distinguish three different regimes in population dynamics that result in different outcomes for the predator-prey interaction (Fig. SI.4).

**Fig. 2.**
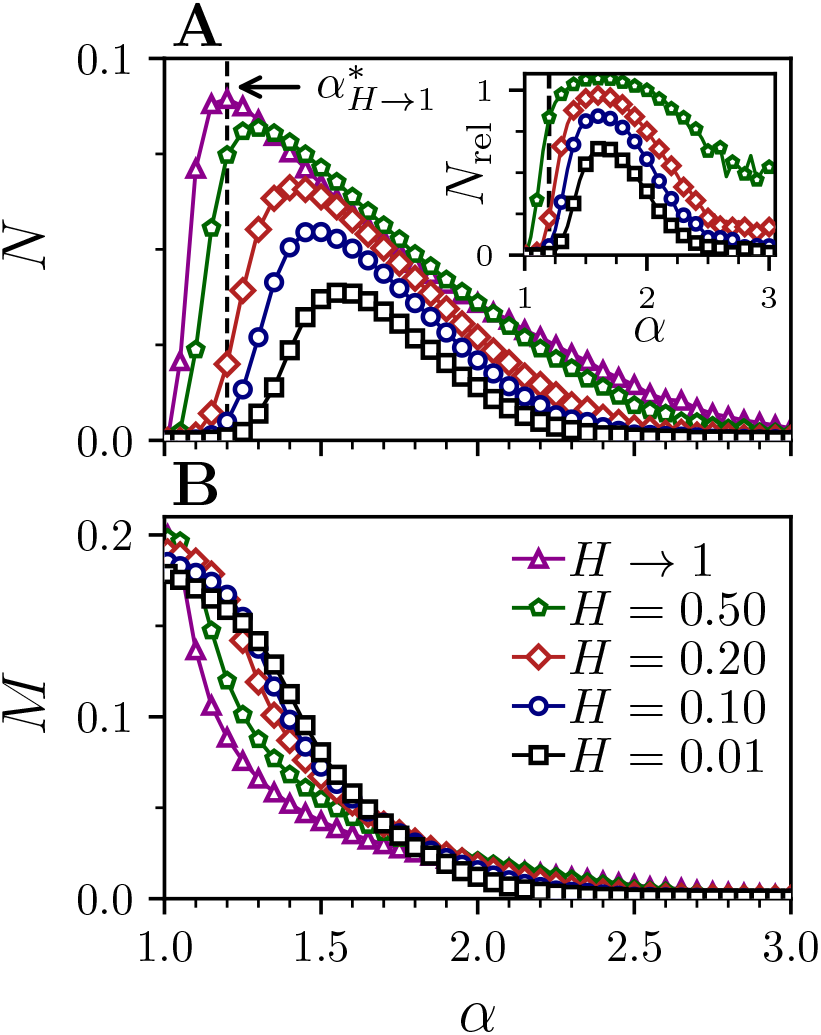
Effect of the Lévy exponent *α* on population densities for different Hurst exponents *H*. Other rate parameters are *μ* = 1/*L, σ* = 0.1 and 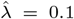 (see SI). (A) Predator density *N*. Dashed vertical line shows optimal Lévy exponent 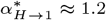 for *H* → 1 and indicates predator extinction if predators cannot rapidly adapt to significant increases in fragmentation (see text). (inset) Relative predator densities *N*_rel_ = *N_H_/N*_*H*→1_ displays decreases in *N* when predators forage with the same *α* in landscapes with higher fragmentation. Note that for some ranges of *α* there exists a preferred intermediate spatial correlation *H* (see text). (B) Prey density *M*. Prey density declines as predators are less dispersive for higher *α*.

First, due to our choice for the predator-prey interaction probability 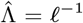, ballistic predators (*α* → 1) rarely consume prey and thus go extinct. Upon predator extinction, prey proliferate until they reach their maximum population size. Notice, however, that this population size does not correspond with the prey habitat density *ρ* in fragmented landscapes because small habitat patches become irreversibly uninhabited (see below).

Second, in the Brownian limit, *α* → 3, predation is intense and prey are overexploited regardless of the level of landscape fragmentation. Interestingly, Brownian-like dispersal effectively induces patch-fidelity and area-restricted (or area-concentrated) search patterns (47–49), as only those foragers that initially spawn near a fragment have the opportunity to reproduce (see SI for more details). This cross-generational patch-fidelity, or predator aggregation, results in prey extinction followed by predator extinction due to lack of prey. Note that predator extinctions are asymptotic due to our choice of the predator death rate and we still observe few individuals in our simulations when they are stopped (see SI).

Third, for intermediate values of the Lévy exponent, our model predicts stable species coexistence at different population sizes that are jointly determined by predator movement, *α*, and habitat fragmentation, *H*. For landscapes that display little fragmentation (*H* → 1), habitat patches are large and predator relocations intersect with prey often. As a result, predation still occurs during the non-reactive phases – represented by long displacements – and predators maximize population densities with near ballistic foraging for *α* ≈ 1.2. In contrast, for highly fragmented landscapes (*H* = 0.01), the model tradeoff between displacement length and prey detection probability becomes more important because predator-prey encounters are more rare. It is thus more critical that predators adopt strategies that increase predation rates while ensuring sufficient encounters with prey. This balance is attained when short displacements are more frequently interspersed with long-range relocations, leading to maximum predator population sizes for *α* ≈ 1.6.

We also find that the range of foraging strategies, i.e., values of *α*, that ensures predator survival becomes more narrow as habitat fragmentation increases (inset Fig. 2A). This result suggests a stronger selective pressure on the foraging strategy in highly fragmented landscapes. Moreover, our results further indicate that foraging strategies that maximize predator population sizes in slightly fragmented landscapes 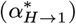 lead to predator extinction as fragmentation increases (Fig. 2A), suggesting a large impact of fragmentation on foraging strategies that result in stable coexistence in weakly fragmented landscapes.

In the intermediate *α* regime, our model suggests that habitat fragmentation does not necessarily affect population densities negatively. Predator populations with *α* < 2 display maximal densities for intermediate values of *H* (Fig. 3), although densities do not significantly decrease when fragmentation decreases (*H* increases). Prey populations can benefit from high levels of fragmentation when predator movement approaches the ballistic regime, approximately for *α* ≤ 2 (Fig. 3B). This benefit results from ballistic predators displaying low prey interaction rates that allow prey to avoid predation by taking advantage of fragmentation and spreading thinly. This aligns with established results obtained for predators performing area-restricted searches (47, 50–52). Such predators exert pressure on prey species to live well spaced-out, and our results indicate that this is also true when predators are Lévy searchers. In our model, however, because prey are sessile and cannot space out during the predator-prey dynamics, they need to adopt such spatial configuration in the initial condition. Moreover, these spatial distributions of prey are more prone to localized extinctions due to stronger demographic fluctuations in small prey patches and to overpredation. This further inhibits the effectiveness of area-restricted search under our model assumptions, as both predator and prey tend to go extinct (Fig. 2, Fig. 3, and see SI).

**Fig. 3.**
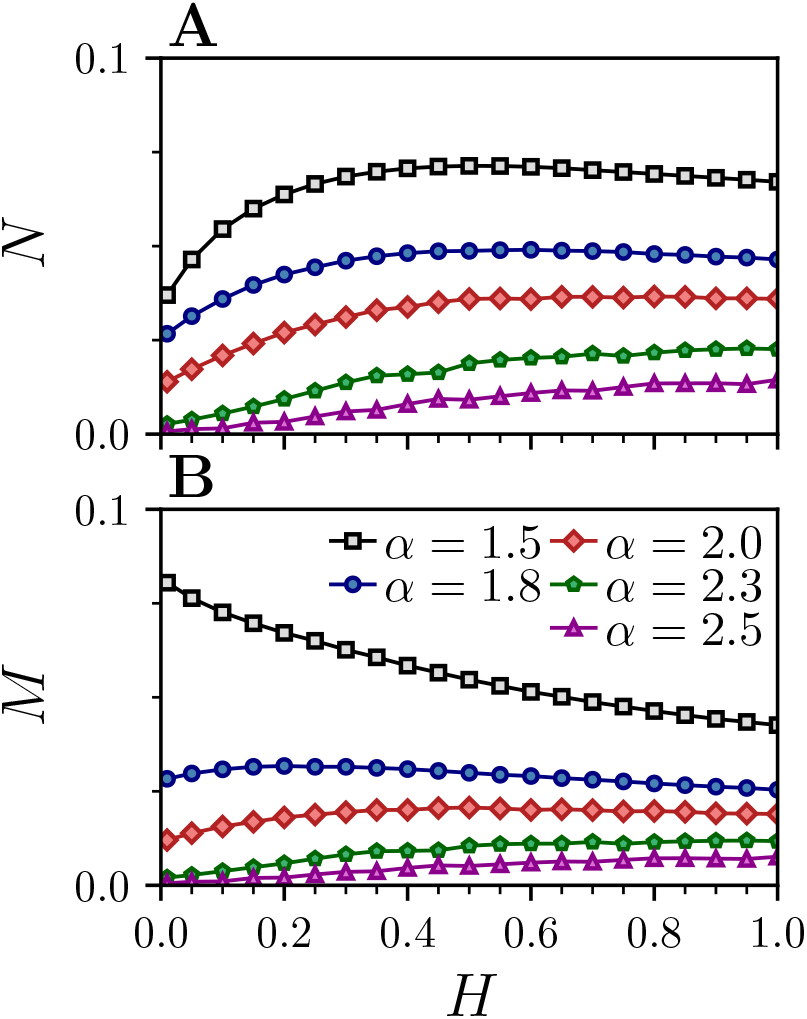
Effect of the Hurst exponent *H* on population densities for different Lévy exponents *α*. Note that, for *α* → 1, we have *N* → 0 as prey encounter rates fall since 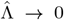. Additionally, for approx. *α* ≥ 3, we have *M* → 0 due to overconsumption. As a result, these values for *α* are not shown. (A) Predator density *N*. For sufficiently high dispersal rates, we observe maximized predator densities for intermediate fragmentation. (B) Prey density *M*. Note that for sufficiently high dispersal rates (low *α*) prey densities are highest in highly fragmented landscapes with *H* → 0 (see text).

Next, we determine ecosystem health using a weighted species richness 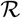 that captures how numerous predator and prey are relative to each other as well as the total population size within the environment (see SI). We define the species richness as

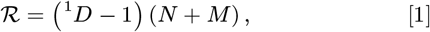

where 1 ≤ ^1^*D* ≤ *S* is the entropy-based diversity index, and *S* = 2 is the total number of species in the system (see see SI and, e.g., Ref. 53). This metric for species richness predominantly follows predator density (Fig. SI.5). However, due to the effect of prey density, the predator foraging strategy that maximizes species richness is consistently more ballistic than that maximizing predator density 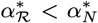 (inset Fig. 6A, Fig. SI.5). This results from lower predator-prey interaction rates 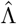 when *α* decreases, which consequently increases prey population size.

### Fragmentation induces irreversible habitat loss

As mentioned above, predators may induce irreversible prey habitat loss in fragmented landscapes because they overexploit small patches that cannot be recolonized because prey individuals are sessile. As a result, following predator extinction, prey population density does not converge to habitat density *ρ* (Fig. 2A). To investigate this further, we measure the patch depletion probability, *P_d_*, as a function of patch size and predator foraging strategy (see SI). Our results indicate that small patches have a higher depletion probability regardless of the predator foraging strategy *α* (Fig. 4), because they host smaller prey populations that are more likely to be exhausted and become subjected to stronger demographic fluctuations. The effect of *α* on the depletion probability is stronger for intermediate patch sizes as higher values of *α* lead to more local predation and, as a consequence, higher patch depletion probability (Fig. 4). Importantly, significant patch depletion occurs even when predators adopt foraging strategies that maximize species richness (Fig. SI.6, Fig. SI.7).

**Fig. 4.**
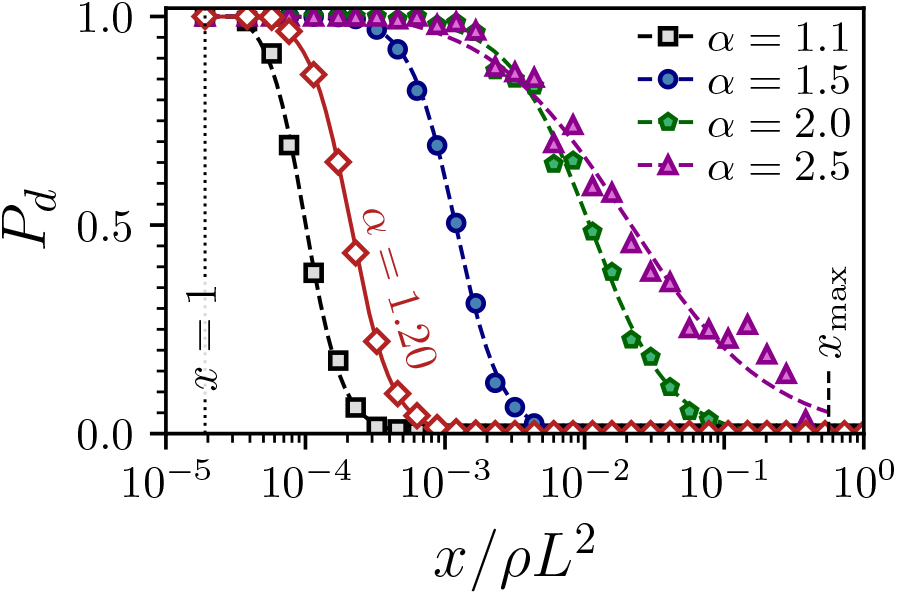
Influence of the Lévy exponent *α* on the probability of patch depletion, *P_d_*, as a function of patch size *x* for intermediate degree of fragmentation with *H* = 0.5. Dotted vertical lines indicate minimum patch sizes at *x* =1 (single site). Dashed vertical line indicates maximum patch size for this particular level of fragmentation and patches with *x* > *x*_max_ do not exist, hence *P_d_*(*x* > *x*_max_) = 0. All other parameters are as in Fig. 2. Solid lines are a guide to the eye. Red curve displays *P_d_* for the optimal response to maximize species richness (*α* = 1.2, see inset Fig. 6). Note that less diffusive foraging strategies (small *α*) result in less depletion as *ρ*_eff_ remains high.

To further evaluate the impact of patch depletion probability *P_d_* on habitat loss, we define the effective habitat density *ρ*_eff_ as the fraction of initial habitat *ρ* that remains available to prey in the quasi-stationary stable state (Fig. 5). Ballistic foragers result in low levels of habitat loss, because predators rapidly go extinct and only a few small patches are depleted (Fig. 4). When *α* increases and short predator displacements become more frequent, the depletion probability is higher for a broader range of patch sizes (compare, for example, the curves for *α* = 1.1 and *α* = 1.5 in Fig. 4). As a result, effective habitat density is a monotonically decreasing function of the Lévy exponent and Brownian foragers minimize the effective habitat density regardless of the level of fragmentation (Fig. 5). However, how much habitat is lost in already fragmented landscapes will depend on the level of fragmentation. For example, Brownian foragers in slightly fragmented landscapes (*H* → 1) eliminate approximately 40% of the initial habitat. In highly fragmented habitats, this percentage is approximately 90% and most of the prey-predator dynamics occurs in the few, (relatively) large patches that remain available for prey.

**Fig. 5.**
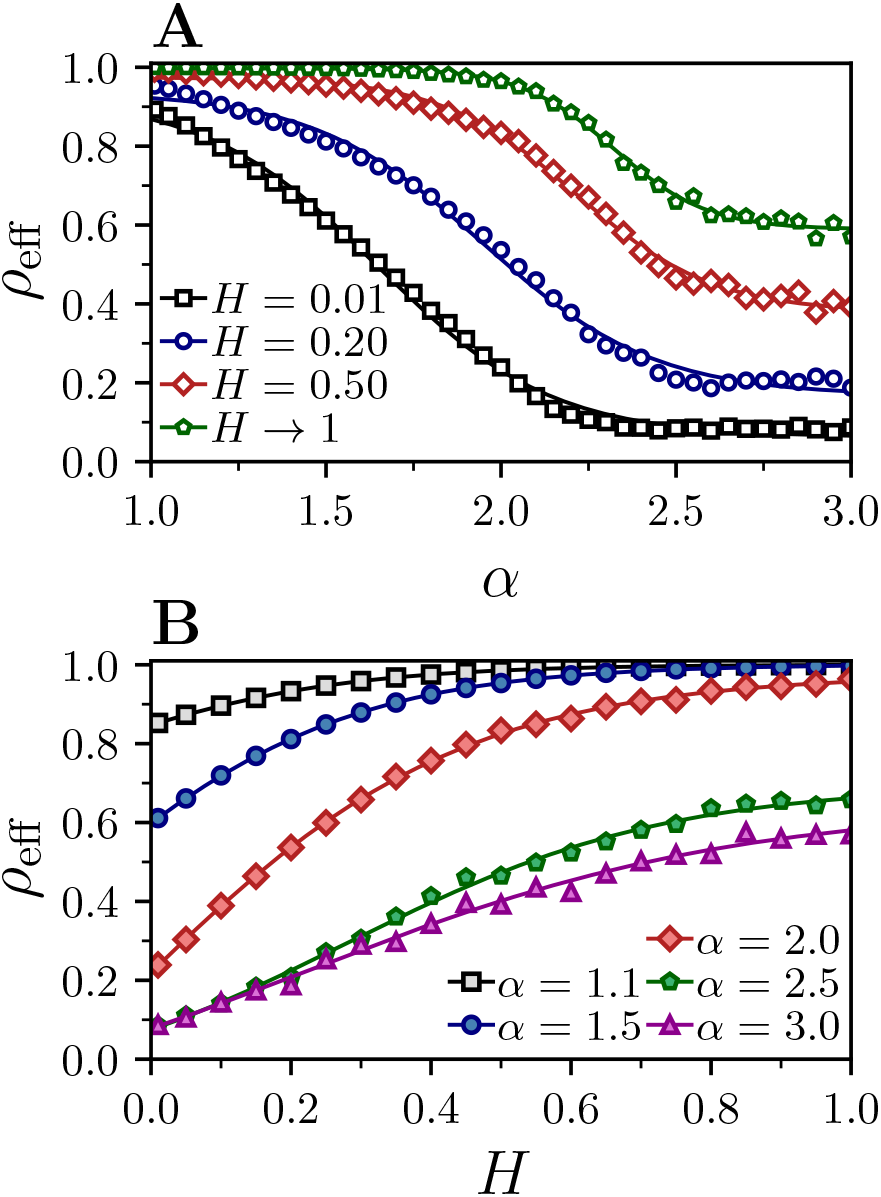
Influence of Lévy exponent *α* and Hurst exponent *H* on the effective habitat density *ρ*_eff_ in the quasi-stationary stable state. All other parameters are as in Fig. 2. Solid lines are a guide to the eye. See also Fig. SI.7 for typical spatial configurations. (A) *ρ*_eff_ as a function of *α* for different *H*, and (B) as a function of the *H* for different *α*.

Finally, we measure how habitat fragmentation affects effective habitat loss for different foraging strategies, *α*. As expected, ballistic predators minimize effective habitat loss because they minimize predation rates (Fig. 5). In contrast, Brownian predators maximize effective habitat loss because they overexploit prey patches locally. Intermediate values of *α* maximize the difference between effective habitat loss at low and high fragmentation (Fig. 5B). For foraging strategies that maximize species richness and predator densities (Fig. 6), increased fragmentation may result in an effective habitat loss of 40%. Importantly effective habitat loss is a nonlinear function of the fragmentation level with much faster decay when landscapes transition from slightly to highly fragmented (Fig. 6A). Population sizes, however, decay much slower in response to increased fragmentation (Fig. 6B), illustrating the importance of foraging strategies in maintaining the stability of ecological communities in response to increased fragmentation and habitat loss.

**Fig. 6.**
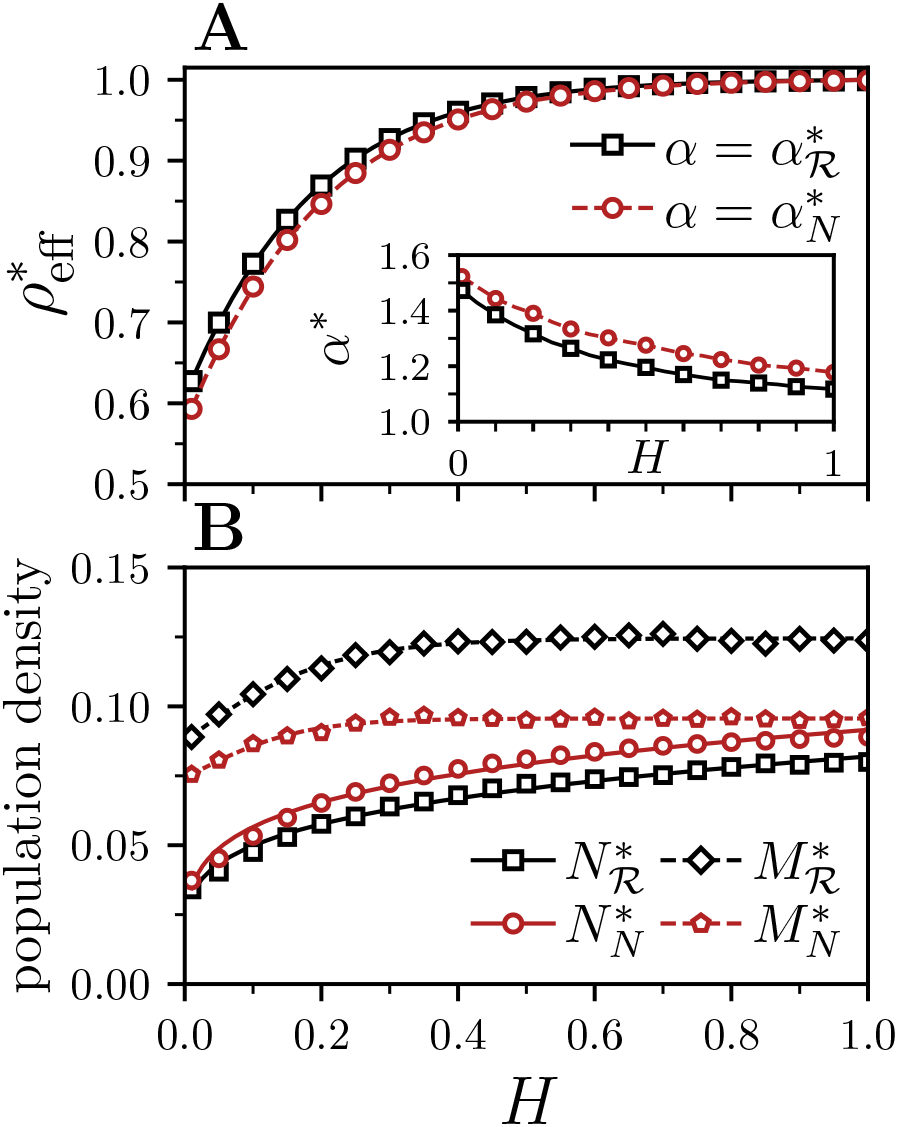
Effect of optimal predator response *α\** to a particular level of fragmentation defined by the Hurst exponent *H*. Optimal responses are either considered to optimize predator population *N* or species richness R, denoted by their respective subscripts. All other parameters are as in Fig. 2. Solid lines are a guide to the eye. (A) Effective habitat density *ρ*_eff_. (inset) Optimal response *α*\*. (B) Predator and prey densities *N* and *M*.

## Discussion

Our stochastic predator-prey model reveals that the interplay between predator foraging behavior and fragmentation strongly influences species persistence, ecosystem stability, and prey habitat conservation (Fig. 7). Predator and prey populations, and the resulting species richness, are maximal for a specific predator foraging strategy *α* that depends nonlinearly on the spatial correlation of habitat *H*. Moreover, increased fragmentation reduces the range of possible *α*-values that result in stable species coexistence (Fig. 2, Fig. 7), which suggests a stronger evolutionary pressure on foraging strategies in highly fragmented environments. We considered here power-law dispersal kernels to model predator foraging, however, similar results have been found for predators exhibiting exponential dispersal kernels (54), which further supports that dispersal is a critical component of long-term ecosystem stability (13, 55). Moreover, as habitat fragmentation increases, prey habitat consists of more and smaller patches. Extinctions within these smaller patches are more likely due to overpredation and stronger demographic fluctuations, ultimately resulting in irreversible prey habitat loss. Our results suggest that optimal predator responses can decrease, but not prevent nor minimize, the amount of lost habitat, and that this reduction is more pronounced when habitat is highly fragmented. The possible outcomes of our model for different predator foraging strategies and habitat fragmentation are summarized in Fig. 7.

**Fig. 7.**
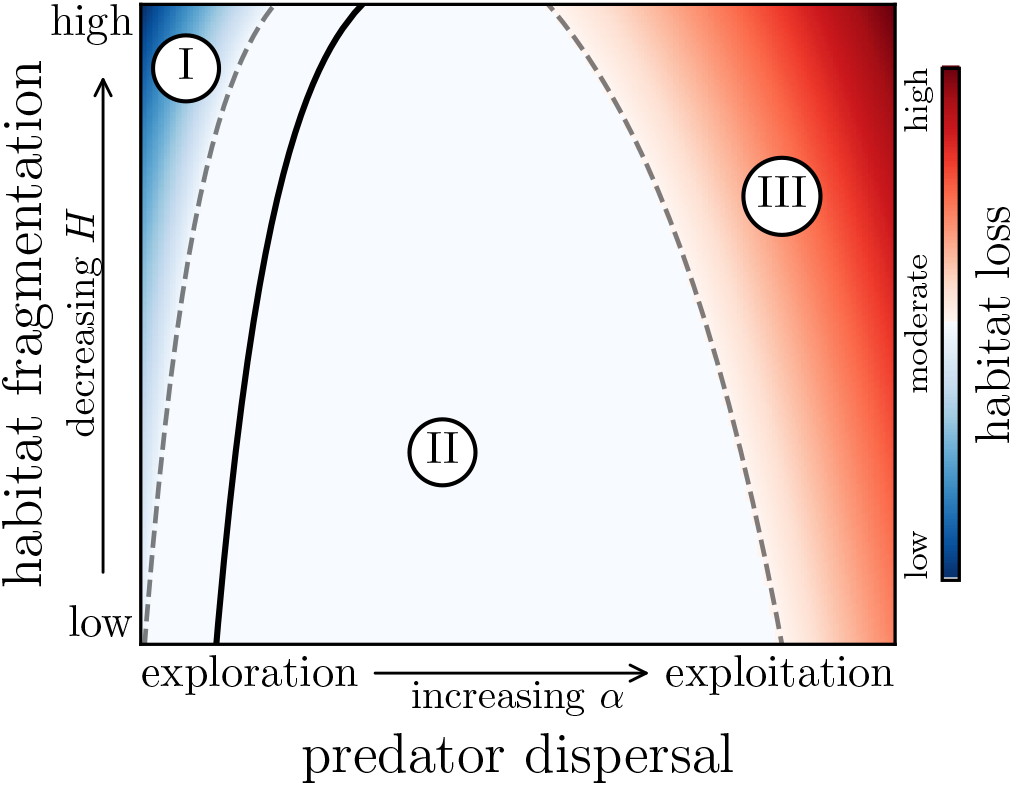
Schematic phase diagram that illustrates model outcomes for population dynamics and irreversible habitat loss as a function of habitat fragmentation and predator dispersal. Our model identifies three qualitatively distinct phases: (I) low habitat loss with predator extinction caused by lack of predation (blue), (II) predator-prey coexistence with moderate habitat loss (white), and (III) high habitat loss due to overexploitation of prey species, leading to full species extinction (red). The black solid line indicates qualitative behavior of optimal dispersal as a function of habitat fragmentation (see also inset Fig. 6). Dashed lines indicate transitions between predator-prey coexistence (II) and predator extinction (I) or full extinction (III). Both the optima and the transitions are gathered and interpolated from simulation data (Fig. 2).

Effective habitat loss mainly results from small patches becoming irreversibly depleted and only large patches remaining inhabitable (Fig. 4). As a result, the effective spatial correlation in the landscape increases, and prey fragmentation decreases resulting in an effective *H* larger than the one used to generate the habitat landscape. Assuming predators can rapidly respond to such a change in fragmentation, possible predator adaptations to this new habitat configuration should result in predators foraging more ballistically (lower *α*, inset Fig. 6). Because habitat loss is less severe for lower values of *α* (Fig. 5), such a response can inhibit further habitat destruction. Hence, our results agree with previous work that indicated that predator dispersal can stabilize irreversible habitat loss and population declines (39, 54, 56–59).

Our model predicts that higher dispersal rates, represented by lower values of *α*, tend to increase ecosystem stability by allowing predators to exploit several prey patches. We neglect, however, all types of dispersal costs that could increase predator death rates when they travel between prey patches (60). Including such costs might be especially relevant when studying the impact of fragmentation on species with low dispersal abilities, such as small mammals (61) and amphibians (62) (but see Ref. 63). Additionally, we also make simplifying assumptions about the landscape. More specifically, we considered a binary lattice and globally fixed demographic rates. Instead, including matrix and edge effects on both predator dispersal and prey reproduction – e.g., by studying a non-binary, heterogeneous habitat matrix (64), movement responses to habitat edges (65), etc. – might reveal potential (de)stabilizing effects that we did not encounter. Furthermore, we did not consider interventions that increase landscape connectivity, e.g., designing corridors to connect spatially separated fragments allowing prey populations to repopulate previously exhausted patches (66). Finally, we omitted habitat heterogeneity in predator reproduction rates, e.g. when different habitat is needed for reproduction, such as aquatic breeding grounds for terrestrial amphibians (67, 68). Future work should incorporate these ingredients and investigate their effects on ecosystem stability.

We also did not investigate possible responses of prey population to predation and habitat loss. In our model, prey is sessile and can only diffuse due to reproduction onto adjacent sites. Therefore, their only response to avoid local extinctions is to increase their reproduction rate *σ*. Hence, environments that contain static prey that cannot cross hard boundaries are subjected to an evolutionary pressure that might favor prey species with higher reproduction rates (69, 70).

We also neglected several features that might affect predator foraging behavior, such as satiation (71), interactions with conspecifics (72, 73), spatial memory (74, 75), long range perception (76, 77), and complex patterns of adaptive movement in response to prey encounters (22). We further considered a trade-off between displacement length and predation efficiency that biases selection against ballistic movement. Although this choice is based on existing literature on intermittent search (27), considering different shapes for this trade-off might alter our results. All these factors can change the optimal foraging strategy in landscapes with varying degrees of fragmentation and hence affect the impact of fragmentation on ecosystem stability.

Finally, higher-order interactions are known to stabilize population dynamics of multi-species systems (78–80) and extinctions in one trophic level may destabilize species’ coexistence and cause extinctions higher up the trophic network (81, 82). Extending our framework to describe multi-species systems with more complex trophic interactions is needed to understand how foraging behavior and fragmentation jointly determine ecosystem stability.

Our work displays the intricate interplay between foraging behavior and habitat fragmentation, and highlights the role of dispersal on population persistence and ecosystem stability in fragmented landscapes. It furthermore shows how increased levels of fragmentation lead to higher irreversible habitat loss and how optimal foraging responses can reduce, but not prevent nor minimize, the amount of lost habitat. Due to their profound ecological consequences, our results suggest that future models of population dynamics should explicitly include optimal foraging arguments when discussing potential effects of landscape fragmentation on ecosystem stability.

## Data accessibility

Python code to reproduce the results of our model is freely available via Zenodo: https://doi.org/10.5281/zenodo.6585088

## Authors’ contributions

J.N.: conceptualization, formal analysis, investigation, methodology, software, data analysis, visualization, writing–original draft, review and editing; P.S.: supervision, writing–review and editing; Y.K.: supervision, writing–review and editing; R.M.G.: conceptualization, investigation, methodology, data analysis, writing–original draft, review and editing;

## Conflict of interest declaration

The authors declare no conflict of interest.

## Funding statement

We received no specific funding for this study.

## Acknowledgments

R.M.G. was supported by FAPESP through grant ICTP-SAIFR 2016/01343-7, Programa Jovens Pesquisadores em Centros Emergentes grants 2019/24433-0 and 2019/05523-8; Instituto Serrapilheira through grant Serra-1911-31200; and the Simons Foundation. The authors would like to thank V. Sudbrack for helpful discussions on generating fragmented landscapes.

## Supplementary Information

### 1. Neutral landscape models for modeling fragmented resource landscapes

Fragmented prey habitat in our model is generated using neutral landscape models (1). More specifically, we generate lattices with periodic boundary conditions from an underlying fractional Brownian motion (fBm) characterized by the Hurst exponent *H* ∈ (0,1) that controls the spatial correlation between adjacent sites (2). We generate lattices of two-dimensional fBm using spectral synthesis (see, e.g., 2–4). As these methods rely on Fourier transforms, they result in periodic lattices with lateral lengths 2^*k*^, with *k* some positive integer. For *H* → 0, adjacent sites are negatively correlated, resulting in landscapes with several small fragments. In contrast, *H* → 1 indicates high correlations between adjacent sites and thus results in landscapes with few, large habitat patches (see Fig. **??**). Because we are interested in landscapes with a specific habitat density *ρ*, we transform the fBm lattice into a binary lattice. As the fBm lattice of size *L* = 2^*k*^ contains 2^*k*^ × 2^*k*^ data points with random values, we select the *ρL*^2^ sites with the highest numerical values to belong to the inhabitable fragments while the remaining sites belong to the uninhabitable matrix (2). These type of landscape models have been used to study landscape connectivity and its effects on animal dispersal (e.g., Refs. 5–8). Note that, during our simulations, the lattice defining a fragmented habitat is static and does not change over time, yet the spatial configuration of prey on this lattice can change due to demographic events (see Stochastic lattice Lotka-Volterra model).

**Fig. SI.1.**
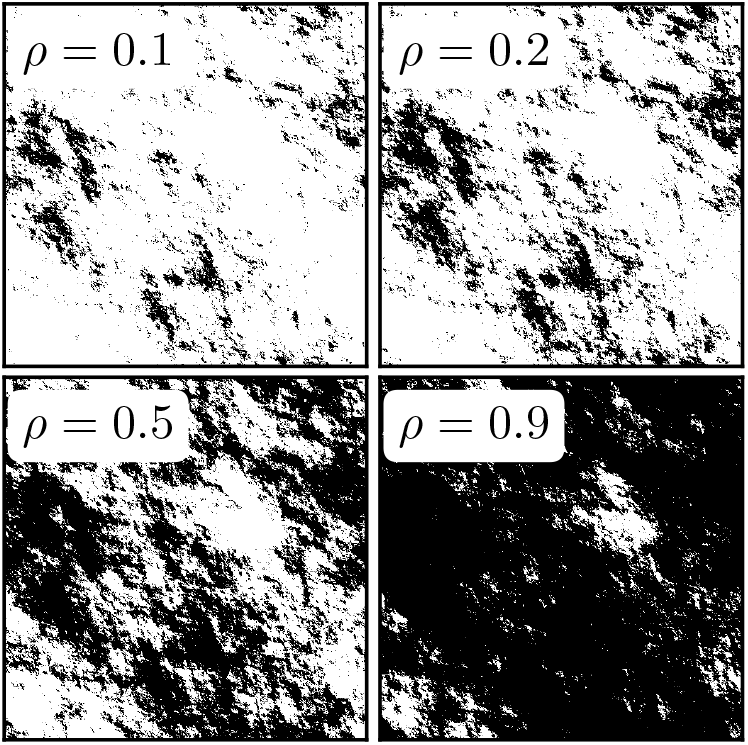
Fragmented landscapes of different prey habitat densities *ρ* for a fixed spatial correlation *H* = 0.1. The effect of fragmentation reduces as *ρ* increases (Fig. **??**).

**Fig. SI.2.**
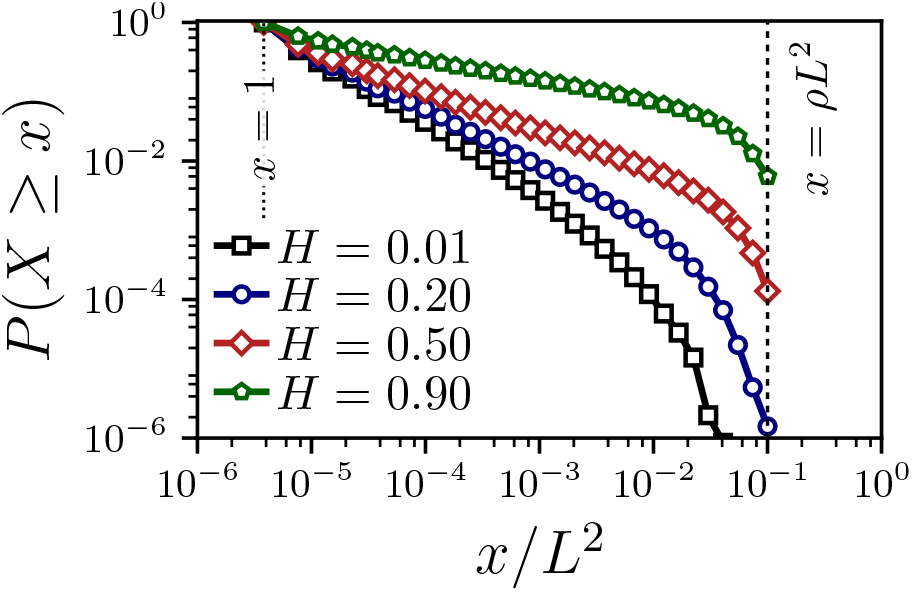
Influence of Hurst exponent *H* on the patch size distribution for *ρ* = 0.2. *P*(*X* ≥ *x*) is the complementary cumulative distribution function of the stochastic variable *X* (i.e., the patch size) being equal or larger than *x*. Results are obtained for 512 × 512 landscapes. The vertical dashed (dotted) line indicates maximum (minimum) possible patch size.

Fragmentation effects are weaker when habitat density is high (Fig. SI.1). This is further exemplified by noting that increased fragmentation decreases average patch sizes, because smaller patches become more frequent as *H* decreases and large patches become (near) nonexistent. (Fig. SI.2). We measure landscape connectivity in terms of the (percolation) probability *p* that a habitat patch connects to either side of the lattice. Because we study systems with periodic boundary conditions, *p* gives the probability of there being a single spanning cluster. For large *ρ* we find that *p* → 1 regardless of the value of *H* (Fig. **??**). This means that the largest patches contain most of the available habitat, which effectively reduces our predator-prey model to one without prey habitat restrictions (i.e., a homogeneous landscape, as in, e.g., Refs. 9, 10).

Because we are interested in the effects of optimal foraging behavior, we study landscapes with habitat density *ρ* = 0.2 for which percolation theory predicts disconnected patch structures for all values of *H* (see, e.g., Ref. 5) and predator dispersal becomes a critical driver of population dynamics. Moreover, as Lévy foragers maximize foraging efficiency only when resource (prey, thus habitat) densities are low (11), only in landscapes with *ρ* (relatively) small would one expect foragers to diffuse anomalously.

### 2. Stochastic lattice Lotka-Volterra model

We study a stochastic lattice Lotka-Volterra model (SLLVM) wherein prey habitat is restricted as a consequence of fragmentation, as we assume they cannot inhabit any site belonging to the hostile matrix. Prey is assumed to be sessile, whereas predators are allowed to disperse freely and can thus move between prey habitat patches. Each lattice site can be in one of four possible states: empty 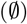, occupied by a predator (*X*), occupied by a prey (*Y*), or occupied by a prey and a predator (*XY*). Multiple occupation by two individuals of the same species (i.e., *XX* or *YY*) is forbidden. The state transitions that fully define the stochastic dynamics of the model are:

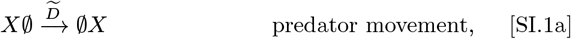

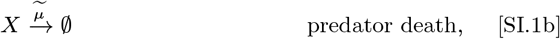

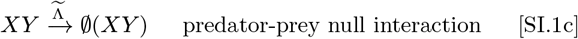

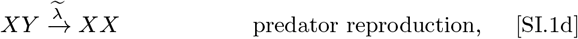

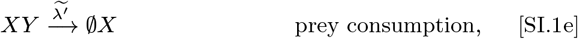

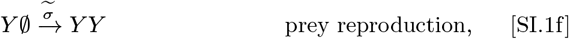

where 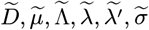, the predator dispersal (diffusion) rate, predator death rate, predator-prey interaction rate, predator reproduction rate, predation rate and the prey reproduction rate, respectively. Note that predator death represents a singlesite reaction, whereas all other processes describe nearest-neighbor two-site reactions.

The above set of stochastic state transitions describes the time evolution of a predator-prey system with demographic fluctuations. We define the spatially averaged predator and prey densities *N* and *M* as the number of predators and prey per unit area, i.e., *N* = *n/L*^2^ and *M* = *m/L*^2^ where *n* and *m* are the number of predators and prey in the lattice of size *L* = 2^*k*^ with *L*^2^ sites, respectively. The system has three possible stationary states: predator and prey extinction, predator extinction followed by prey proliferation, and stable predator-prey coexistence. First, the zero-abundance fixed point (*N, M*) = (0, 0) is reached when both predator and prey have gone extinct. This fixed point results from overconsumption of prey by predators, which leads to prey and subsequently predator extinction. Second, the prey-proliferation fixed point (*N, M*) = (0, *ρ_M_*) results from systems wherein only predators have gone extinct. This is an effect of underconsumption of prey that leads to predator extinction, followed by prey proliferating on the available habitat due to the absence of predation. Note, however, that following predator extinction prey densities *ρ_M_* might not reach the initial habitat density *ρ* due to overpredation, thus local extinction, in the smallest prey patches (i.e., *ρ_M_* < *ρ*, see also Fig. **??**). The final stable coexistence fixed points with *N,M* > 0 are those where predators neither over- nor underconsume (Fig. SI.4).

Although our SLLVM is similar to existing models, (e.g, Refs. 9, 12), it differs in two main ingredients. First, we allow for a null predator-prey interaction in which prey consumption is not followed by predator reproduction. This additional model ingredient is included in our model because initial empirical simulation results reported that truncation effects (see Numerical implementation) resulted in non-Lévy predator dispersal. As such, the null-interaction ensures that predator dispersal displays similar (scale-free) characteristics as optimal forager movement in sparse resource landscapes. Second, as we consider predators that perform Lévy walks, we introduce a previously unexplored explicit spatiotemporal coupling. More specifically, whereas displacement in Ref. 12 was instantaneous due to predators following Lévy flights, predators in our model can at most displace a single lattice unit per step. Here, a step refers to the event that a single lattice site that contains a predator is randomly chosen (see Numerical implementation). As a result, the duration of each relocation event is proportional to the length of the (sampled) displacement. For more details regarding the differences between Lévy walks and Lévy flights, we refer the interested reader to detailed descriptions on this topic, such as Ref. 13.

### 3. Numerical implementation

Below, we provide details on the numerical implementation of the SLLVM and discuss the choice of specific values for the demographic rates.

#### A. Monte Carlo simulations of the restricted SLLVM

We consider a Monte Carlo approach for simulating the stochastic dynamics defined by the state transitions of Eq. (SI.1). A single Monte Carlo time step corresponds to selecting all occupied lattice sites once on average. In our Monte Carlo algorithm, we randomly select occupied sites and update their state according to the following rules:

##### Monte Carlo update rules for the SLLVM

- (*predator death*) If the selected site contains a predator, it dies with probability *μ*;
- If the predator survived;
  – (*predator dispersal*) if the adjacent site is empty, move there and continue the current relocation (see below);
  – (*relocation truncation*) if the adjacent site contains a predator, truncate the current relocation and do not move;
  – if the adjacent site contains prey, either
    * (*double occupancy*) move there, but do *not* interact with the prey with probability Λ;
    * (*predator reproduction*) truncate the current relocation, and reproduce by adding a predator that replaces the prey with probability *λ*;
    * (*prey consumption*) truncate the current relocation, and consume prey with probability *λ′* = 1 – Λ – *λ*. Upon prey consumption (i.e., no reproduction) the selected site is emptied and the prey is replaced with a predator;
- (*prey reproduction*) If the selected site contains a prey individual, choose a habitable adjacent site randomly (if any) and, if the chosen site is empty, place a prey there with probability *σ*.

Here, *μ*, Λ, *λ* and *σ* are the probabilities for predator death, predator-prey interaction, predator reproduction and prey reproduction respectively. Note that we have *λ* + *λ’* + Λ = 1, thus prey consumption *without* predator reproduction occurs with probability *λ*^′^. Because the predator-prey interaction probability Λ depends on the length of predator displacement (see below), it is convenient to introduce the following relations:

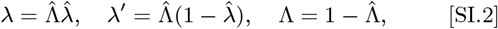

where 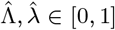 the conditional probabilities of the corresponding state transitions of our model. More specifically, 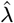 is the conditional predator reproduction probability given that it interacts with prey. Predator-prey interaction occurs with probability 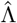. For example, consider 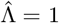, thus 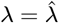 and hence 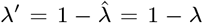, i.e., 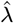 is the probability of prey consumption with predator reproduction and 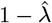 the probability of prey consumption without reproduction. In contrast, when predators never interact with prey for 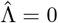, there is no reproduction nor consumption, as we find 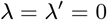.

#### B. Sampling of Lévy walks on a lattice

Predators perform Lévy walks on a lattice. Each straight line displacement has a length *ℓ* sampled from a discrete power law distribution with exponent *α*, i.e., *p*(*ℓ*) ∝ *ℓ^-α^*. To avoid integer overflows when generating samples from this distribution when movement becomes ballistic (*α* → 1), we truncated the power law such that *ℓ* ∈ [*I*_min_, *ℓ*_max_] (14). We set the lower truncation at the unit lattice spacing *ℓ*_min_ = 1 and choose the upper truncation as to ensure that long flights result in low predator-prey interaction probability, i.e., 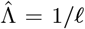, cf. intermittent random searches (see main text and, e.g., Ref. 15). More specifically, we choose *ℓ*_max_ such that the predator-prey interaction probability is, in practice, negligible when predators perform ballistic motion with *α* → 1. As we take averages over a number of independent model realizations in the order 10^2^–10^3^, we choose *ℓ*_max_ such that 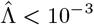 when *α* → 1. Because predator-prey interaction is a Bernoulli trial, with the average number of predator-prey encounters for a ballistic predator equal to *ρL*, *ℓ*_max_ ~ 10^5^ ensures that 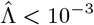. Note however, that it is very unlikely to observe such long predator displacements in our simulations. The reason is that the probability of a predator surviving long enough to complete such a long displacement is extremely small as the mortality rate *μ* = 1/*L* (see below). As a result, ballistic foragers, in practice, do not interact with prey for *ℓ*_max_ ~ 10^5^, as desired by having 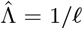 (see also Fig. **??**). Specifically, we choose *ℓ*_max_ = 200*L* = 102400, as *L* = 512, in all of our experiments.

Next, we define the discrete random variable 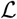 that has a truncated power law distribution *p*(*ℓ*) ∝ *ℓ*^-*α*^ for *ℓ* ∈ [*ℓ*_min_, *ℓ*_max_]. To generate random samples that follow this distribution, we first define the discrete complementary cumulative distribution *P*(*ℓ*) (the survival function) as the probability of 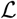 to be larger than some length *ℓ* (14, 16):

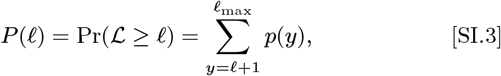

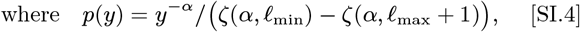

with 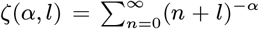 the Hurwitz-*ζ* function. To sample predator displacement lengths, we draw a random number *r* from a uniform distribution between 0 and 1 and compute *ℓ* that satisfies *P*(*ℓ*) = 1 – *r*. As *P*(*ℓ*) cannot be inverted in closed form, we execute a binary search within the interval [*ℓ*_min_, *ℓ*_max_] to solve for *ℓ* (14, 16). Because we are interested in discrete samples, we continue the binary search until the value of *ℓ* is narrowed down to *k* ≤ *ℓ* < *k* + 1, for some integer *k*. Then, we discard the non-integer part of *ℓ* to be used as the discrete sample. Binary search is implemented in many standard libraries and can be executed efficiently. Even so, one profits from pre-computing the Hurwitz-*ζ* functions for all values *ℓ* ∈ [*ℓ*_min_, *ℓ*_max_], as computation of these values can be (relatively) computationally expensive.

**Fig. SI.3.**
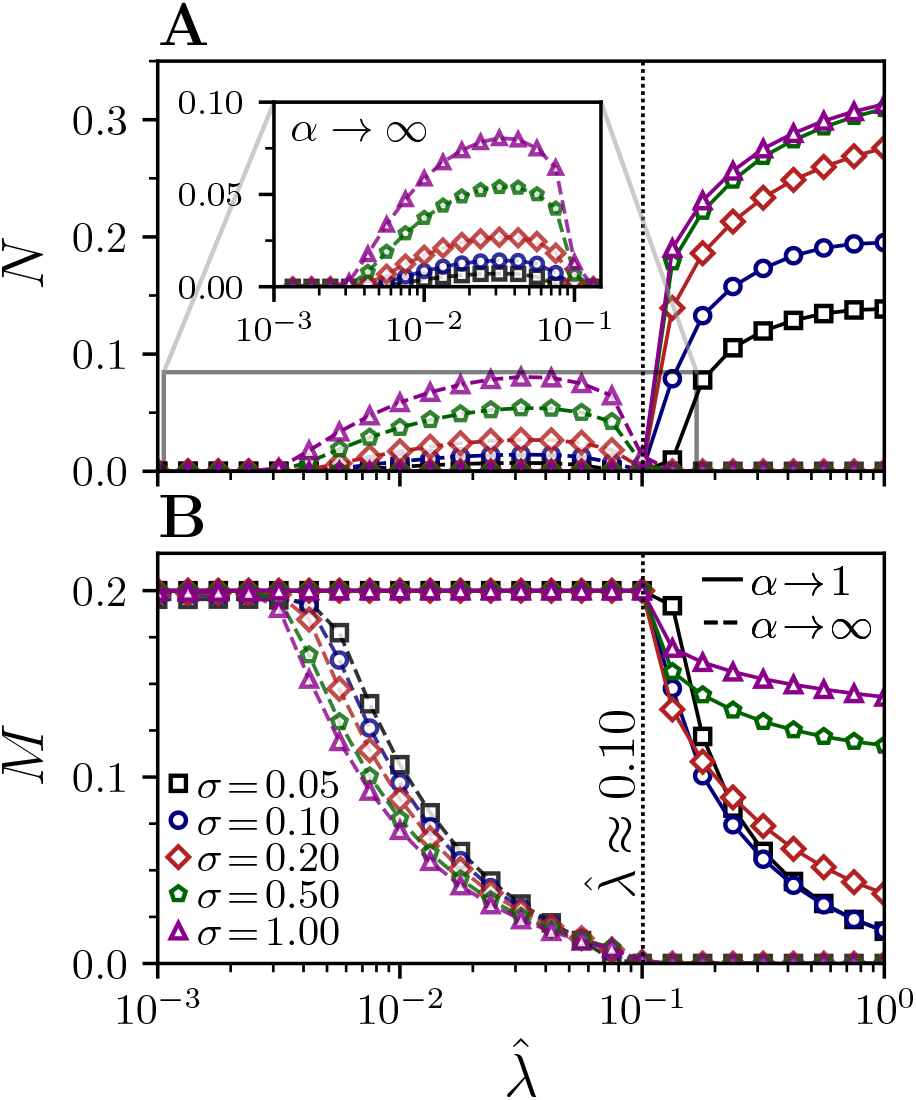
Influence of prey reproduction probability *σ* on population densities in the quasi-stationary stable state versus the predator reproduction probability 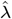. Predator reproduction probability is given by 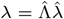 with predator-prey interaction probability 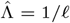 and predator death probability *μ* = 1/*L*. To determine demographic rates for which the landscape is most fragile, we compare ballistic predators (solid lines, *α* → 1, Λ → 1) with predators that do nearest neighbor random walks (dashed lines, *α* → ∞, Λ = 0), cf. Ref. 12. Environments are generated with *ρ* = 0.2 and little to no fragmentation with *H* → 1 (see Fig. **??**). (A) Predator density *N*. Inset displays more detail on predator densities for predators with *α* → ∞. (B) Prey density *M*. Vertical dotted lines in (A) and (B) at *λ* ≈ 0.1 indicate the predator reproduction rate for which the ecosystem is most fragile.

#### C. Ecosystem fragility and choice of demographic rates

Since the dynamics of population densities depend critically on the demographic rates, we select specific values to represent Lévy predators in fragile ecosystems as to maximize the impact of fragmentation. More specifically, because we let predators follow Lévy walks, the probability of predator death *μ* should be such that displacement lengths are not exponentially truncated due to predator death. We therefore fix a low predator death rate, *μ* =1/*L*, that allows them to make long system-length displacements within their lifetime. Next, we choose predator and prey reproduction rates such that systems with little fragmentation (*H* → 1) are most fragile. Here, fragile systems are those wherein markedly different predator dispersal rates result in predator extinction either through overconsumption (nearest neighbor random walks for *α* → ∞) or through underconsumption (ballistic motion for *α* → 1) (12). Our results indicate that, regardless of the prey reproduction probability *σ*, systems are most fragile for predator reproduction probability 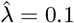 (Fig. SI.3). Note that this specific parameter choice represents ecologically relevant scenarios in which predators are highly motile, long-lived, and reproduce slowly. Finally, we choose *σ* = 0.1 for the prey reproduction rate. An overview of the used parameters and their specific values is listed in Table 1. Typical population dynamics obtained using this parameterization are shown in Fig. SI.4.

**Table 1.**
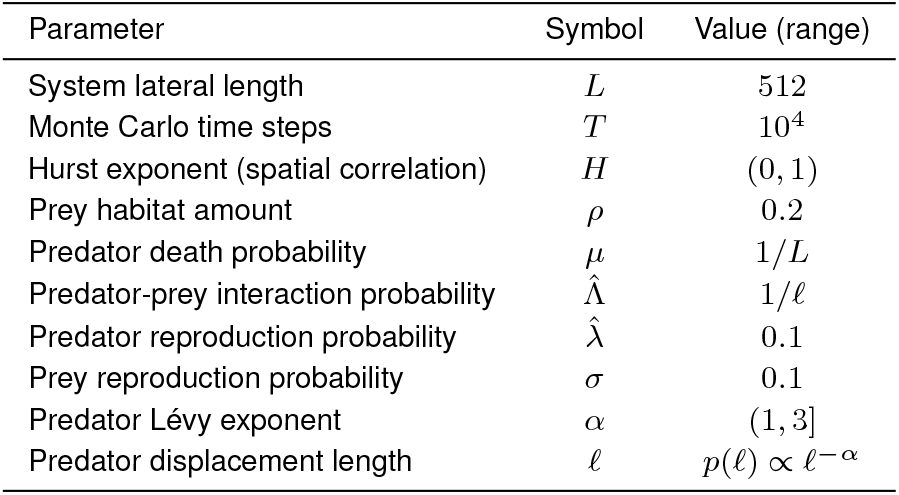
Overview of model parameterization.

**Fig. SI.4.**
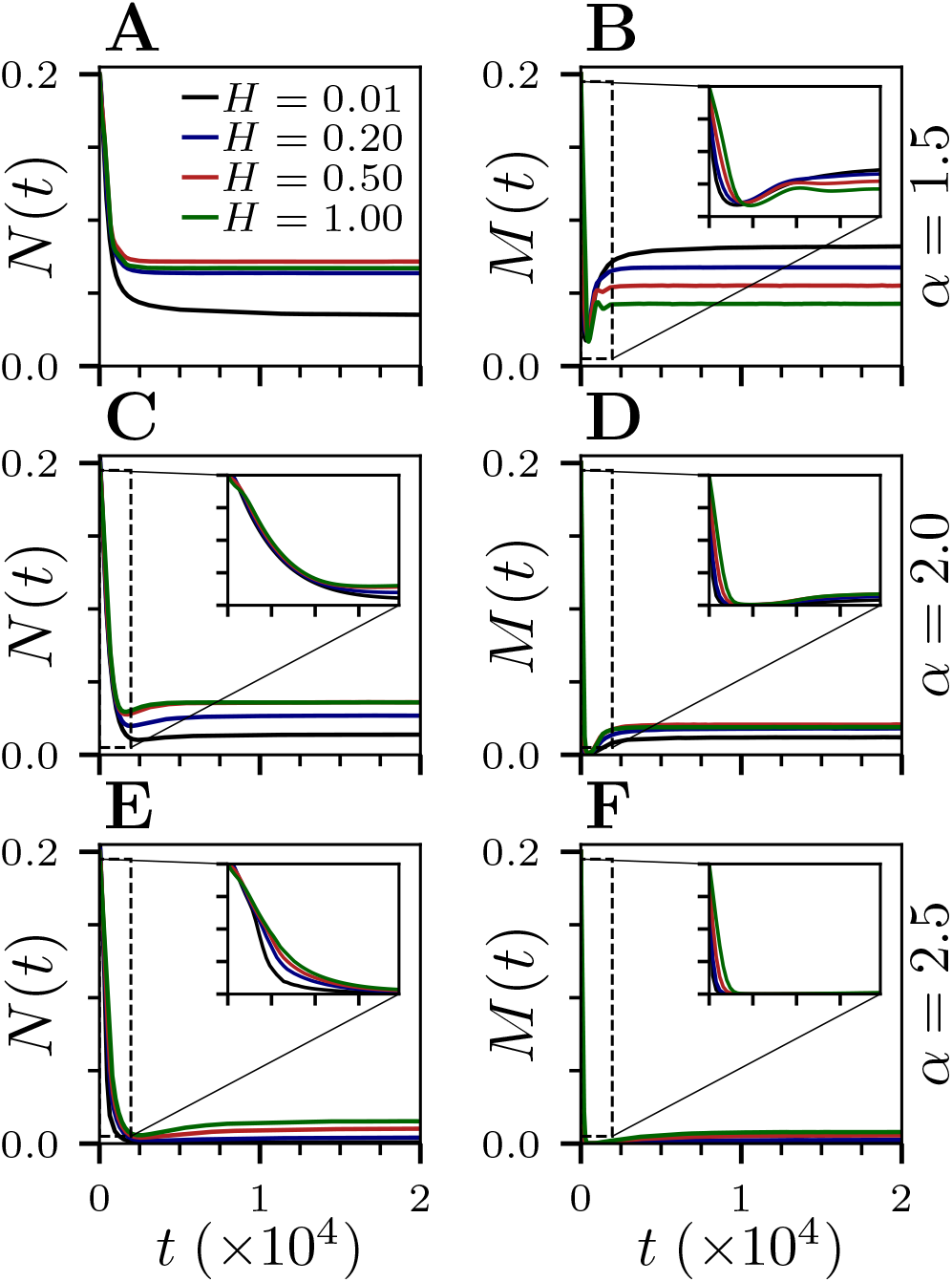
Time evolution of population densities for several Lévy exponents and Hurst exponents *H* and the number of Monte Carlo time steps *T* = 2 · 10^4^. Insets show short-term dynamics up to *t* = 2 · 10^3^ Monte Carlo time steps. Note how quasi-stationary stable states are attained for *t* ≈ 10^4^, which is the number of Monte Carlo time steps used in all other simulations. For *α* → 1 and *α* ≥ 3 we find predator extinction and prey extinction (followed by predator extinction) respectively (not shown, but see Fig. **??**). All other parameters are as in Fig. **??**.

### 4. Species richness and ecosystem health

While population densities are important to determine ecosystem stability, it can be worthwhile to additionally discuss ecosystem ‘health’. In ecology, diversity indices are often used to indicate ecosystem health, where primers such as biodiversity (e.g., the number of different species) are often of interest (17). In particular, *species richness* measures the number of species relative to the total number of individuals. We consider the effective species’ diversity, Hill number, or ‘true diversity’, ^*q*^*D*, to capture ecosystem health (see e.g., Ref. 18 for an overview). The effective species’ diversity is an entropy-based measure and defined as

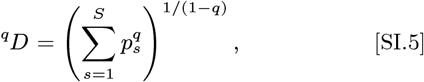

where *S* is the total number of species, and *p_s_* the probability of sampling species *s* when sampling randomly from the total population of all individuals, i.e., the proportional abundance of species *s*. Here, *q* defines the sensitivity of the true diversity to species’ relative abundance, where *q* > 1 weighs the more abundant species more heavily and *q* < 1 weighs rare species more heavily. For *q* =1, all species are equally weighed and the true diversity can be defined by using the limit 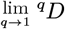:

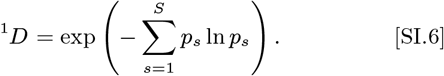

**Fig. SI.5.**
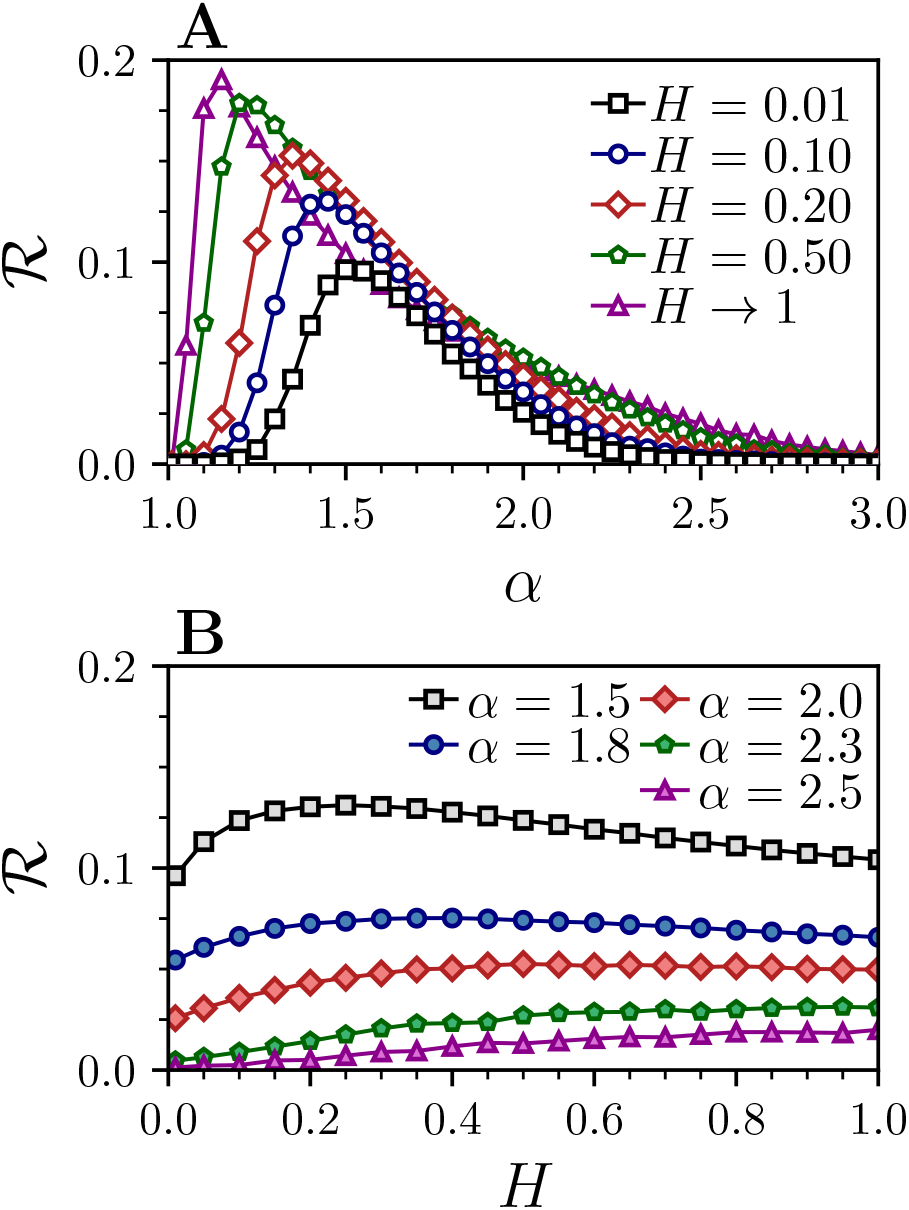
Influence of Lévy exponent *α* and Hurst exponent *H* on the species richness 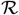. Note that 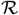 mainly follows predator density (Fig. **??**, Fig. **??**), but effects of prey density results in consistently more ballistic dispersal to maximize 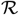 when compared to predator density. All other parameters are as in Fig. **??**.

However, the diversity index with *q* = 1 does not take into account the total population size and is effectively biased towards systems with equal number of species. For example, consider a two species system – as the predator-prey system studied here – with *N* predators and *M* prey individuals. Such a system is as ‘diverse’ for *N* = *M* = 1 as for *N*′ = *M*′ = 100, while we consider the second system more ‘healthy’, simply because it contains more organisms. To correct for this bias, we define the weighted species richness as 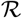 as

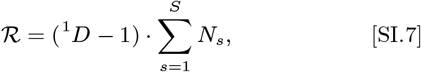

where *N_s_* is the spatial average of the population density of species *s* per unit area (see Stochastic lattice Lotka-Volterra model). We subtract 1 from the true diversity to bring its value between 0 and 1. Briefly, 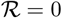 corresponds to systems where there is only one species remaining, whereas 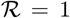 represents situations in which both species persist with equal population sizes. The species richness 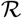 as a function of *α* and *H* is shown in Fig. SI.5. Note that other definitions of species richness exist and that their definitions should depend strongly on the context wherein it is applied (cf. Ref. 19). For our intended purposes, i.e., a two species predator-prey model, the definitions in Eqs. (SI.6), (SI.7) serve as a simple indicator of the biodiversity – thus ecosystem health – in different model realizations. We did not observe changes in maxima (i.e., optimal responses) to different entropy-based measures of ecosystem health.

### 5. Patch depletion probability

We compute the patch depletion probability *P_d_* as a function of patch size *x* by checking if the separated patches (fragments) on the lattice contain prey at the end of our simulations. If they do not contain prey, we consider them depleted and compute the depletion probability of patches of a specific size *x* over a number of independent model realizations. Patch depletion probabilities depend strongly on both predator dispersal and habitat fragmentation. Here, we would like to emphasize that, when landscapes display high levels of fragmentation, patches of smaller sizes are more frequent. This, together with increased patch depletion probabilities for small patches (Fig. **??**), increases habitat loss in highly fragmented systems (Fig. **??**, Fig. **??**). More importantly, patch depletion probabilities are considerable even when predator’s foraging patterns maximize species richness 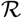 (Fig. SI.6, Fig. SI.7).

**Fig. SI.6.**
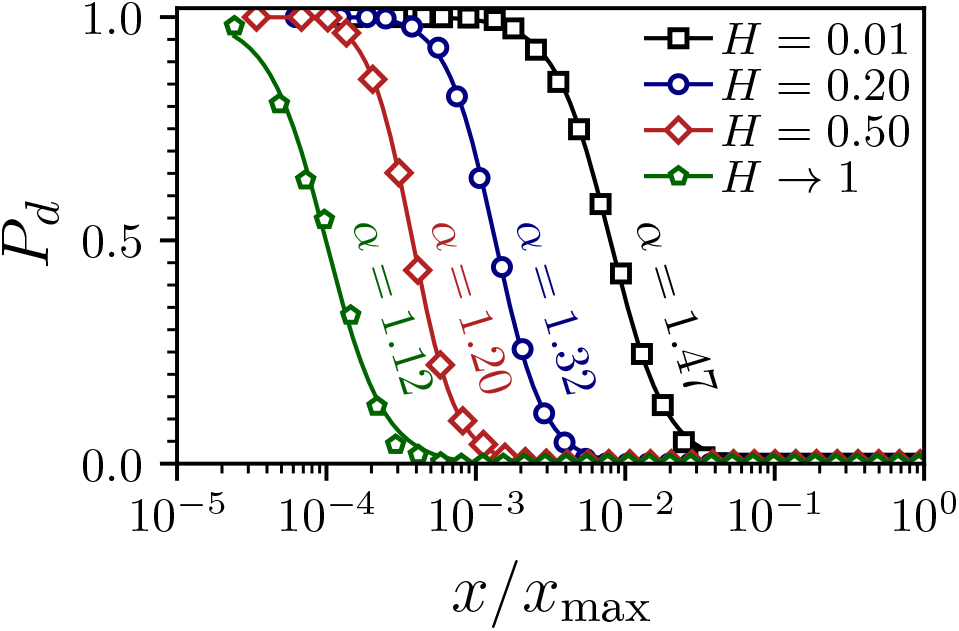
Influence of Hurst exponent *H* on the probability of patch depletion, *P_d_*, as a function of patch size *x* for Lévy exponents that maximize species richness 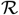 (i.e., 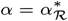, as extracted from Fig. **??**). Note the normalization by *x*_m_ax as landscapes with low *H* contain smaller patches (Fig. **??**).

**Fig. SI.7.**
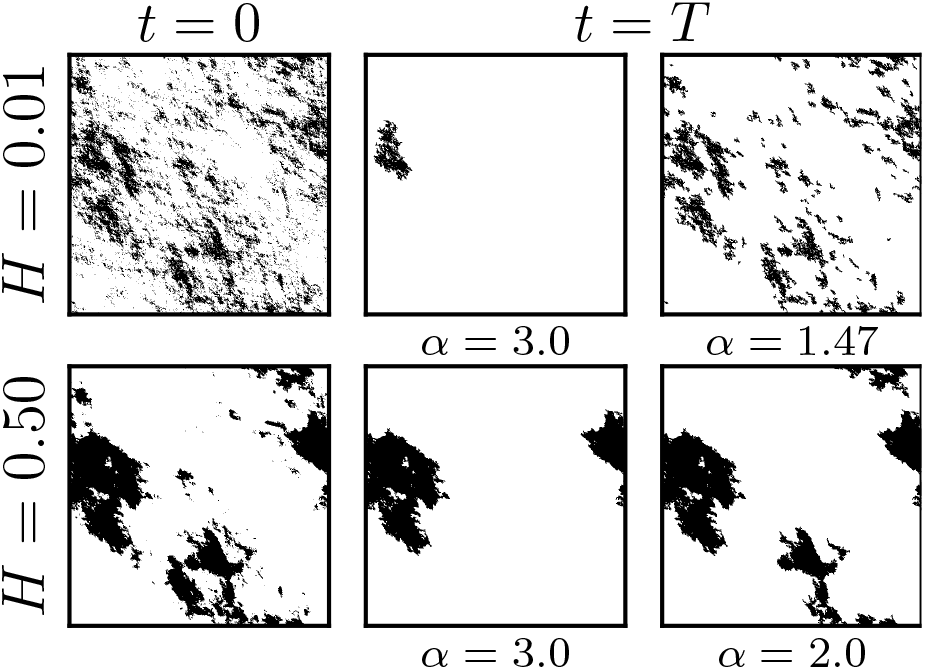
Typical example of the effects of fragmentation and foraging strategy on habitat loss and effective habitat. Black and white regions depict prey habitat and matrix, respectively. Left most panels depict initial habitat at *t* = 0. Other panels depict the system at the end of our simulations *t* = *T* = 10^4^. (top) Habitat loss in a highly fragmented environment. For Brownian predators (*α* = 3.0) nearly all habitat is lost, while optimal predator foraging (*α* = 1.47) greatly reduces, but not prevents nor minimizes, habitat loss. Note that full extinction (i.e. with *ρ*_eff_ = 0) can also occur. (bottom) Habitat loss in an environment with intermediate fragmentation. Brownian predators deplete a large patch, while scale-free predators (*α* = 2.0) preserve a large habitat amount. Note that the optimal predator foraging strategy (*α* ≈ 1.2 for *H* = 0.50) does not result in considerable habitat loss (Fig. **??**), and is therefore not shown.

### 6. Lévy walks, patch-fidelity and area-restricted searches

The biological relevance of Lévy walks as a foraging strategy is currently a subject of debate (20–22). In particular, area-restricted searches have been shown to outperform Lévy walks in environments wherein resources are clumped (23–26). In area-restricted searches, predators switch between exploitative or exploratory behavior based on sensory input, such as the number of recent prey encounters. The sensitivity with which predators modulate their behavior is hereby of critical importance, and is usually assumed to increased search time in areas of high prey density and decrease traveling time through low prey density areas (27–30). It should be noted, however, that area-restricted searches do not necessarily result in full patch depletion (i.e. overexploitation), as foragers can switch from exploitation to exploration when there are still resources present in the patch. In our model, the cross-generational patch-fidelity that occurs for large *α* should not be confused with area-restricted searches, as predators do not change *α* depending on sensory input. If this is the case, and when prey is assumed to be sessile, our results indicate that fragmentation greatly increases the frequency of local prey extinction events. We find that this effect is exacerbated when predators display patch-fidelity, which occurs for large values of *α* (Fig. **??**, Fig. **??**). Therefore, patch-fidelity does not favor species’ coexistence and ecosystem stability because predators do not properly balance patch exploitation and new patch exploration in a way that allows prey to recover locally (12). As a result, our model suggests that, analogously, area-restricted searches that lead to full patch depletion are detrimental for ecosystem stability. Moreover, area-restricted search behavior might actually decrease system resilience to fragmentation. Future work should investigate how area-restricted searches influence population dynamics in fragmented systems and extensions of our model that consider more complex adaptive patterns of movement, e.g. based on inter- and intraspecific interactions, could shed more light on this issue.

## Notes

### Competing Interest Statement

The authors have declared no competing interest.

### Summary of Updates

Slight adaptations of text to increase clarity in correspondence with peer review process

## References

1. NM Haddad, et al., Habitat fragmentation and its lasting impact on earth’s ecosystems. Sci. advances 1, e1500052 (2015).

2. JM Lord, DA Norton, Scale and the spatial concept of fragmentation. Conserv. Biol. 4, 197–202 (1990).

3. AB Franklin, BR Noon, TL George, What is habitat fragmentation? Stud. avian biology 25, 20–29 (2002).

4. KR Crooks, et al., Quantification of habitat fragmentation reveals extinction risk in terrestrial mammals. Proc. national Acad. Sci. 114, 7635–7640 (2017).

5. L Fahrig, Ecological responses to habitat fragmentation per se. Annu. Rev. Ecol. Evol. Syst. 48, 1–23 (2017).

6. RJ Fletcher Jr, et al., Is habitat fragmentation good for biodiversity? Biol. conservation 226, 9–15 (2018).

7. PO Cheptou, AL Hargreaves, D Bonte, H Jacquemyn, Adaptation to fragmentation: evolutionary dynamics driven by human influences. Philos. Transactions Royal Soc. B: Biol. Sci. 372, 20160037 (2017).

8. L Fahrig, Effects of habitat fragmentation on biodiversity. Annu. review ecology, evolution, systematics 34, 487–515 (2003).

9. P Kareiva, Population dynamics in spatially complex environments: theory and data. Philos. Transactions Royal Soc. London. Ser. B: Biol. Sci. 330, 175–190 (1990).

10. RA Briers, Incorporating connectivity into reserve selection procedures. Biol. conservation 103, 77–83 (2002).

11. JC Williams, CS ReVelle, SA Levin, Spatial attributes and reserve design models: a review. Environ. Model. & Assess. 10, 163–181 (2005).

12. PR Armsworth, JE Roughgarden, The impact of directed versus random movement on population dynamics and biodiversity patterns. The Am. Nat. 165, 449–465 (2005).

13. BB Niebuhr, et al., Survival in patchy landscapes: the interplay between dispersal, habitat loss and fragmentation. Sci. reports 5, 1–10 (2015).

14. A Beckerman, OL Petchey, PJ Morin, Adaptive foragers and community ecology: linking individuals to communities and ecosystems. Funct. Ecol. 24, 1–6 (2010).

15. HH Wei, F Lutscher, From individual movement rules to population level patterns: the case of central-place foragers in Dispersal, Individual Movement and Spatial Ecology. (Springer), pp. 159–175 (2013).

16. MA Lewis, PK Maini, SV Petrovskii, Dispersal, individual movement and spatial ecology. Lect. Notes Math. (Mathematics Biosci. Series) 2071 (2013).

17. W Campeau, AM Simons, B Stevens, The evolutionary maintenance of lévy flight foraging. PLOS Comput. Biol. 18, e1009490 (2022).

18. GM Viswanathan, MG Da Luz, EP Raposo, HE Stanley, The physics of foraging: an introduction to random searches and biological encounters. (Cambridge University Press), (2011).

19. V Zaburdaev, S Denisov, J Klafter, Lévy walks. Rev. Mod. Phys. 87, 483 (2015).

20. GM Viswanathan, et al., Lévy flight search patterns of wandering albatrosses. Nature 381, 413–415 (1996).

21. AM Reynolds, et al., Displaced honey bees perform optimal scale-free search flights. Ecology 88, 1955–1961 (2007).

22. NE Humphries, et al., Environmental context explains Lévy and Brownian movement patterns of marine predators. Nature 465, 1066–1069 (2010).

23. AM Foley, et al., Purposeful wanderings: mate search strategies of male white-tailed deer. J. Mammal. 96, 279–286 (2015).

24. LR Paiva, et al., Scale-free movement patterns in termites emerge from social interactions and preferential attachments. Proc. Natl. Acad. Sci. 118 (2021).

25. S Benhamou, How many animals really do the lévy walk? Ecology 88, 1962–1969 (2007).

26. M Plank, A James, Optimal foraging: Lévy pattern or process? J. The Royal Soc. Interface 5, 1077–1086 (2008).

27. O Bénichou, C Loverdo, M Moreau, R Voituriez, Intermittent search strategies. Rev. Mod. Phys. 83, 81 (2011).

28. GM Viswanathan, et al., Optimizing the success of random searches. Nature 401, 911 (1999).

29. F Bartumeus, J Catalan, UL Fulco, ML Lyra, GM Viswanathan, Optimizing the encounter rate in biological interactions: Lévy versus Brownian strategies. Phys. Rev. Lett. 88, 097901 (2002).

30. F Bartumeus, MGE da Luz, GM Viswanathan, J Catalan, Animal search strategies: a quantitative random-walk analysis. Ecology 86, 3078–3087 (2005).

31. K Zhao, et al., Optimal Lévy-flight foraging in a finite landscape. J. The Royal Soc. Interface 12, 20141158 (2015).

32. A Ferreira, E Raposo, G Viswanathan, M Da Luz, The influence of the environment on Lévy random search efficiency: fractality and memory effects. Phys. A: Stat. Mech. its Appl. 391, 3234–3246 (2012).

33. ME Wosniack, MC Santos, EP Raposo, GM Viswanathan, MGE da Luz, Robustness of optimal random searches in fragmented environments. Phys. Rev. E 91, 052119 (2015).

34. JMC Hutchinson, PM Waser, Use, misuse and extensions of “ideal gas” models of animal encounter. Biol. Rev. 82, 335–359 (2007).

35. HB Jackson, L Fahrig, What size is a biologically relevant landscape? Landsc. ecology 27, 929–941 (2012).

36. ND Jackson, L Fahrig, Habitat amount, not habitat configuration, best predicts population genetic structure in fragmented landscapes. Landsc. Ecol. 31, 951–968 (2016).

37. JP O’Dwyer, Beyond an ecological ideal gas law. Nat. Ecol. Evol. 4, 14–15 (2020).

38. KJ Duffy, Simulations to investigate animal movement effects on population dynamics. Nat. Resour. Model. 24, 48–60 (2011).

39. T Dannemann, D Boyer, O Miramontes, Lévy flight movements prevent extinctions and maximize population abundances in fragile Lotka-Volterra systems. Proc. Natl. Acad. Sci. 115, 3794–3799 (2018).

40. RH Gardner, BT Milne, MG Turnei, RV O’Neill, Neutral models for the analysis of broad-scale landscape pattern. Landsc. ecology 1, 19–28 (1987).

41. Q Wang, GP Malanson, Neutral landscapes: bases for exploration in landscape ecology. Geogr. Compass 2, 319–339 (2008).

42. DL Kramer, RL McLaughlin, The behavioral ecology of intermittent locomotion. Am. Zool. 41, 137–153 (2001).

43. D Campos, F Bartumeus, V Méndez, Search times with arbitrary detection constraints. Phys. Rev. E 88, 022101 (2013).

44. F Bartumeus, et al., Foraging success under uncertainty: search tradeoffs and optimal space use. Ecol. letters 19, 1299–1313 (2016).

45. RJ Fletcher Jr, Multiple edge effects and their implications in fragmented landscapes. J. Animal Ecol. 74, 342–352 (2005).

46. M Pfeifer, et al., Creation of forest edges has a global impact on forest vertebrates. Nature 551, 187–191 (2017).

47. JA Wiens, Population responses to patchy environments. Annu. review ecology systematics 7, 81–120 (1976).

48. P Kareiva, G Odell, Swarms of predators exhibit “preytaxis” if individual predators use area-restricted search. The Am. Nat. 130, 233–270 (1987).

49. H Weimerskirch, D Pinaud, F Pawlowski, CA Bost, Does prey capture induce area-restricted search? A fine-scale study using GPS in a marine predator, the wandering albatross. The Am. Nat. 170, 734–743 (2007).

50. N Tinbergen, M Impekoven, D Franck, An experiment on spacing-out as a defence against predation. Behaviour 28, 307–320 (1967).

51. J Taylor, The advantage of spacing-out. J. Theor. Biol. 59, 485–490 (1976).

52. DR Formanowicz Jr, MS Bobka, Predation risk and microhabitat preference: an experimental study of the behavioral responses of prey and predator. Am. Midl. Nat. pp. 379–386 (1989).

53. H Tuomisto, A consistent terminology for quantifying species diversity? Yes, it does exist. Oecologia 164, 853–860 (2010).

54. E Colombo, C Anteneodo, Metapopulation dynamics in a complex ecological landscape. Phys. Rev. E 92, 022714 (2015).

55. D Roff, Population stability and the evolution of dispersal in a heterogeneous environment. Oecologia 19, 217–237 (1975).

56. K Johst, R Brandl, S Eber, Metapopulation persistence in dynamic landscapes: the role of dispersal distance. Oikos 98, 263–270 (2002).

57. G Bohrer, R Nathan, S Volis, Effects of long-distance dispersal for metapopulation survival and genetic structure at ecological time and spatial scales. J. Ecol. 93, 1029–1040 (2005).

58. A Trakhtenbrot, R Nathan, G Perry, DM Richardson, The importance of long-distance dispersal in biodiversity conservation. Divers. Distributions 11, 173–181 (2005).

59. A Gupta, T Banerjee, PS Dutta, Increased persistence via asynchrony in oscillating ecological populations with long-range interaction. Phys. Rev. E 96, 042202 (2017).

60. D Bonte, et al., Costs of dispersal. Biol. reviews 87, 290–312 (2012).

61. AS Pires, PK Lira, FA Fernandez, GM Schittini, LC Oliveira, Frequency of movements of small mammals among atlantic coastal forest fragments in brazil. Biol. Conserv. 108, 229–237 (2002).

62. M Alex Smith, D M. Green, Dispersal and the metapopulation paradigm in amphibian ecology and conservation: are all amphibian populations metapopulations? Ecography 28, 110–128 (2005).

63. WC Funk, AE Greene, PS Corn, FW Allendorf, High dispersal in a frog species suggests that it is vulnerable to habitat fragmentation. Biol. letters 1, 13–16 (2005).

64. TH Ricketts, The matrix matters: effective isolation in fragmented landscapes. The Am. Nat. 158, 87–99 (2001).

65. G Andreguetto Maciel, F Lutscher, How individual movement response to habitat edges affects population persistence and spatial spread. Am. Nat. 182, 42–52 (2013).

66. P Kindlmann, F Burel, Connectivity measures: a review. Landsc. ecology 23, 879–890 (2008).

67. CG Becker, CR Fonseca, CFB Haddad, RF Batista, PI Prado, Habitat split and the global decline of amphibians. Science 318, 1775–1777 (2007).

68. CR Fonseca, et al., Modeling habitat split: landscape and life history traits determine amphibian extinction thresholds. PLoS One 8, e66806 (2013).

69. IP Owens, PM Bennett, Ecological basis of extinction risk in birds: habitat loss versus human persecution and introduced predators. Proc. Natl. Acad. Sci. 97, 12144–12148 (2000).

70. BE Sæther, Ø Bakke, Avian life history variation and contribution of demographic traits to the population growth rate. Ecology 81, 642–653 (2000).

71. Y Li, BC Rall, G Kalinkat, Experimental duration and predator satiation levels systematically affect functional response parameters. Oikos 127, 590–598 (2018).

72. R Martínez-García, JM Calabrese, T Mueller, KA Olson, C López, Optimizing the search for resources by sharing information: Mongolian gazelles as a case study. Phys. Rev. Lett. 110, 248106 (2013).

73. R Martínez-García, JM Calabrese, C López, Optimal search in interacting populations: Gaussian jumps versus lévy flights. Phys. Rev. E 89, 032718 (2014).

74. WF Fagan, et al., Spatial memory and animal movement. Ecol. letters 16, 1316–1329 (2013).

75. J Nauta, Y Khaluf, P Simoens, Hybrid foraging in patchy environments using spatial memory. J. Royal Soc. Interface 17, 20200026 (2020).

76. WF Fagan, et al., Perceptual ranges, information gathering, and foraging success in dynamic landscapes. The Am. Nat. 189, 474–489 (2017).

77. R Martinez-Garcia, CH Fleming, R Seppelt, WF Fagan, JM Calabrese, How range residency and long-range perception change encounter rates. J. theoretical biology 498, 110267 (2020).

78. K McCann, A Hastings, GR Huxel, Weak trophic interactions and the balance of nature. Nature 395, 794–798 (1998).

79. A Dobson, et al., Habitat loss, trophic collapse, and the decline of ecosystem services. Ecology 87, 1915–1924 (2006).

80. J Grilli, G Barabás, MJ Michalska-Smith, S Allesina, Higher-order interactions stabilize dynamics in competitive network models. Nature 548, 210–213 (2017).

81. OJ Schmitz, PA Hambäck, AP Beckerman, Trophic cascades in terrestrial systems: a review of the effects of carnivore removals on plants. The Am. Nat. 155, 141–153 (2000).

82. WJ Ripple, et al., What is a trophic cascade? Trends ecology & evolution 31, 842–849 (2016).

## References

1. Robert H Gardner and Dean L Urban. Neutral models for testing landscape hypotheses. Landscape Ecology, 22(1):15–29, 2007.

2. Dietmar Saupe. Algorithms for random fractals. In The science of fractal images, pages 71–136. Springer, 1988.

3. Patrick Flandrin. Wavelet analysis and synthesis of fractional brownian motion. IEEE Transactions on information theory, 38(2):910–917, 1992.

4. Joseph D Chipperfield, Calvin Dytham, and Thomas Hovestadt. An updated algorithm for the generation of neutral landscapes by spectral synthesis. PLoS One, 6(2):e17040, 2011.

5. Kimberly A With, Sean J Cadaret, and Cinda Davis. Movement responses to patch structure in experimental fractal landscapes. Ecology, 80(4):1340–1353, 1999.

6. Thomas Hovestadt, Stefan Messner, and Joachim Poethke Hans. Evolution of reduced dispersal mortality and ‘fat-tailed’dispersal kernels in autocorrelated landscapes. Proceedings of the Royal Society of London. Series B: Biological Sciences, 268(1465):385–391, 2001.

7. Dries Bonte, Thomas Hovestadt, and Hans-Joachim Poethke. Evolution of dispersal polymorphism and local adaptation of dispersal distance in spatially structured landscapes. Oikos, 119(3):560–566, 2010.

8. Nathan D Jackson and Lenore Fahrig. Habitat amount, not habitat configuration, best predicts population genetic structure in fragmented landscapes. Landscape Ecology, 31(5):951–968, 2016.

9. Mauro Mobilia, Ivan T. Georgiev, and Uwe C. Täuber. Fluctuations and correlations in lattice models for predator-prey interaction. Phys. Rev. E, 73:040903, Apr 2006..

10. Ulrich Dobramysl, Mauro Mobilia, Michel Pleimling, and Uwe C Täuber. Stochastic population dynamics in spatially extended predator–prey systems. Journal of Physics A: Mathematical and Theoretical, 51(6):063001, 2018..

11. Gandimohan M Viswanathan, Sergey V Buldyrev, Shlomo Havlin, MGE Da Luz, EP Raposo, and H Eugene Stanley. Optimizing the success of random searches. Nature, 401(6756):911, 1999.

12. Teodoro Dannemann, Denis Boyer, and Octavio Miramontes. Lévy flight movements prevent extinctions and maximize population abundances in fragile Lotka-Volterra systems. Proceedings of the National Academy of Sciences, 115(15):3794–3799, 2018.

13. V Zaburdaev, S Denisov, and J Klafter. Lévy walks. Reviews of Modern Physics, 87(2):483, 2015.

14. Hong Zhu, Yingchao Xie, and Maochao Xu. Discrete truncated power-law distributions. Australian & New Zealand Journal of Statistics, 58(2):197–209, 2016.

15. Olivier Bénichou, Claude Loverdo, Michel Moreau, and Raphael Voituriez. Intermittent search strategies. Reviews of Modern Physics, 83(1):81, 2011.

16. Aaron Clauset, Cosma Rohilla Shalizi, and Mark EJ Newman. Power-law distributions in empirical data. SIAM review, 51(4):661–703, 2009.

17. Robert H Whittaker. Evolution and measurement of species diversity. Taxon, 21(2-3):213–251, 1972.

18. Hanna Tuomisto. A consistent terminology for quantifying species diversity? Yes, it does exist. Oecologia, 164(4):853–860, 2010.

19. Root Gorelick. Commentary: Do we have a consistent terminology for species diversity? The fallacy of true diversity. Oecologia, 167(4):885–888, 2011.

20. Simon Benhamou. How many animals really do the lévy walk? Ecology, 88(8):1962–1969, 2007.

21. Alex James, Michael J Plank, and Andrew M Edwards. Assessing Lévy walks as models of animal foraging. Journal of the Royal Society Interface, 8(62):1233–1247, 2011.

22. Graham H Pyke. Understanding movements of organisms: it’s time to abandon the Lévy foraging hypothesis. Methods in Ecology and Evolution, 6(1):1–16, 2015.

23. AM Reynolds. Adaptive Lévy walks can outperform composite Brownian walks in non-destructive random searching scenarios. Physica A: Statistical Mechanics and its Applications, 388(5):561–564, 2009.

24. AS Ferreira, EP Raposo, GM Viswanathan, and MGE Da Luz. The influence of the environment on Lévy random search efficiency: fractality and memory effects. Physica A: Statistical Mechanics and its Applications, 391(11):3234–3246, 2012.

25. Thomas T Hills, Christopher Kalff, and Jan M Wiener. Adaptive Lévy processes and area-restricted search in human foraging. PLoS One, 8(4):e60488, 2013.

26. Johannes Nauta, Pieter Simoens, and Yara Khaluf. Group size and resource fractality drive multimodal search strategies: A quantitative analysis on group foraging. Physica A: Statistical Mechanics and its Applications, 590:126702, 2022.

27. Peter Kareiva and Garrett Odell. Swarms of predators exhibit “preytaxis” if individual predators use area-restricted search. The American Naturalist, 130(2):233–270, 1987.

28. Peter Turchin. Translating foraging movements in heterogeneous environments into the spatial distribution of foragers. Ecology, 72(4):1253–1266, 1991.

29. Simon Benhamou. Efficiency of area-concentrated searching behaviour in a continuous patchy environment. Journal of Theoretical Biology, 159(1):67–81, 1992.

30. Peter D Walsh. Area-restricted search and the scale dependence of path quality discrimination. Journal of Theoretical Biology, 183(4):351–361, 1996.

